# Structure and mechanism of NALCN-FAM155A-UNC79-UNC80 channel complex

**DOI:** 10.1101/2021.12.20.473568

**Authors:** Yunlu Kang, Lei Chen

**Author notes:** Correspondence and lead contact: Lei Chen.

## Abstract

NALCN channel mediates sodium leak currents and is important for maintaining proper resting membrane potential. NALCN and FAM155A form the core complex of the channel, the activity of which essentially depends on the presence of both UNC79 and UNC80, two auxiliary proteins. NALCN, FAM155A, UNC79, and UNC80 co-assemble into a large hetero-tetrameric channel complex. Genetic mutations of NALCN channel components lead to neurodevelopmental diseases. However, the structure and mechanism of the intact channel complex remain elusive. Here, we present the cryo-EM structure of the mammalian NALCN-FAM155A-UNC79-UNC80 quaternary complex. The structure showed that UNC79-UNC80 form a large piler-shaped heterodimer which was tethered to the intracellular side of the NALCN channel through tripartite interactions with the cytoplasmic loops of NALCN. Two interactions are essential for proper cell surface localization of NALCN. The other interaction relieves the self-inhibition of NALCN by pulling the auto-inhibitory CTD Interacting Helix (CIH) out of its binding site.

## Introduction

NALCN channel mediates voltage-modulated sodium leak currents which can be blocked by extracellular calcium ^1^. It is essential for maintaining the proper resting membrane potential, and thus the electrical excitability of certain cells ^2^. The mutation of the NALCN gene can lead to genetic diseases such as Infantile Hypotonia with Psychomotor Retardation and Characteristic Facies 1 (IHPRF1; OMIM 615419)^3,4^ and Congenital Contractures of the Limbs and Face, Hypotonia, and Developmental Delay (CLIFAHDD; OMIM 616266)^5^. Mice lacking the NALCN gene are neonatal lethal due to respiratory rhythm defects ^6^, while the gain-of function mutation of NALCN impairs rapid eye movement sleep and causes the *dreamless* phenotype in mouse ^7^. Functional NALCN channel is a hetero-tetrameric channelosome that is composed of NALCN, FAM155, UNC79, and UNC80 proteins, with a total molecular weight of 920 kDa ^1^. The co-expression of these four proteins is necessary and sufficient to reconstitute robust NALCN currents in a heterologous system such as Xenopus oocyte and HEK293 cells ^1^. NALCN protein is the pore-forming subunit of the complex and shares sequence and structural homology with eukaryotic voltage-gated sodium channels (Na_V_) and calcium channels (Ca_V_)^8^. FAM155 proteins are transmembrane proteins with a cysteine-rich domain. They are important for the membrane localization of NALCN ^9^. UNC79 and UNC80 are large proteins without any known domains but are essential for the function of NALCN ^10–12^. Notably, the lack of either UNC79 or UNC80 resulted in no NALCN currents in a heterologous system, such as HEK293T cells or Xenopus oocyte ^1^. In agreement with this, genetic mutations of the UNC80 gene also lead to IHPRF diseases (IHPRF2; OMIM 616801) in human ^13^, and knock-out of UNC79 leads to disrupted breathing rhythms ^14^, phenocopying the loss-of-function of the NALCN channel in mice ^6^. Recently advances on the structure determination of mammalian NALCN-FAM155A subcomplex have provided insights into how FAM155A interacts with the NALCN channel, how the functional and degenerate voltage sensors are spatially arranged, and how the non-canonical selectivity filter allows the permeation of sodium ^15–17^ Moreover, the high-resolution structure allowed the identification of a CTD Interacting Helix (CIH) on the linker of NALCN domain II-III (D_II-III_)^15^. The interaction between CTD and CIH has never been observed in related Na_V_ or Ca_V_ channels before. Despite the progress, these structural works were done using the NALCN- FAM155A subcomplex which is essentially not functional due to the absence of UNC79 and UNC80 subunits ^1^. Therefore, the structures of UNC79 and UNC80 and their regulatory mechanism on the NALCN channel remain enigmatic. To answer these fundamental questions, we embarked on structural studies of the NALCN-FAM155A-UNC79-UNC80 hetero-tetrameric channel complex.

## Results and discussion

### Architecture of the NALCN-FAM155A-UNC79-UNC80 quaternary complex

Previous studies on the NALCN-FAM155A heterodimer showed that this subcomplex is well-folded in the absence of UNC79 and UNC80 ^15–17^ However, both UNC79 and UNC80 are crucial for the NALCN currents in HEK293T cells ^1^, indicating that UNC79 and UNC80 might not be necessary for channel folding, but rather affect the plasma membrane localization of the channel or play other regulatory roles. We inserted 3×HA tag after P1077 of NALCN which is on the extracellular loop between S5 and P1 helices of Domain III (D_III_) and found that the insertion of 3×HA tag does not affect electrophysiological properties of NALCN channel (Supplementary information, Fig. S1a). We used HA antibody to quantify the surface localization of NALCN ^18^. Using this construct, we found that the NALCN-FAM155A subcomplex has little surface localization (Fig. 1a), which is consistent with the fact that little currents were observed when only NALCN and FAM155A were expressed ^1^. Co-expression of both UNC79 and UNC80 dramatically promoted the surface localization of NALCN (Fig. 1a), in agreement with markedly enhanced currents ^1^. These results collectively suggest that co-expression of UNC79 and UNC80 could promote the surface localization of the NALCN to enhance NALCN currents. To understand the underlying structural mechanism, we sought to obtain the NALCN-FAM155A-UNC79-UNC80 quaternary protein complex for cryo-EM studies. By screening the high-affinity binary interactions within the complex, we found that UNC79 and UNC80 subunits co-migrate on FSEC (Supplementary information, Fig. S1b), suggesting they could form a stable heterodimer. However, the quaternary complex tends to dissociate during purification ^16,17^, probably due to the low affinity between NALCN-FAM155A subcomplex and UNC79-UNC80 heterodimer. To overcome this obstacle, we exploited the tight binding between GFP and its nanobody ^19^. We fused GFP onto the C-terminus of NALCN and fused nanobody (NbGFP) onto the N-terminus of UNC80 to increase the affinity between NALCN-FAM155A and UNC79-UNC80 and to enhance the stability of the hetero-tetrameric complex (Supplementary information, Fig. S1c). We also included HA or FLAG tags in certain subunits to facilitate the detection during protein expression and purification. We found that co-expression of NALCN-GFP, FAM155A-FLAG, NbGFP-UNC80, and HA-UNC79 conferred typical NALCN currents in HEK293T cells (Supplementary information, Fig. S1d), suggesting these constructs recapitulate the properties of NALCN and are suitable for studying the structural mechanism of the NALCN quaternary complex.

**Fig. 1:**
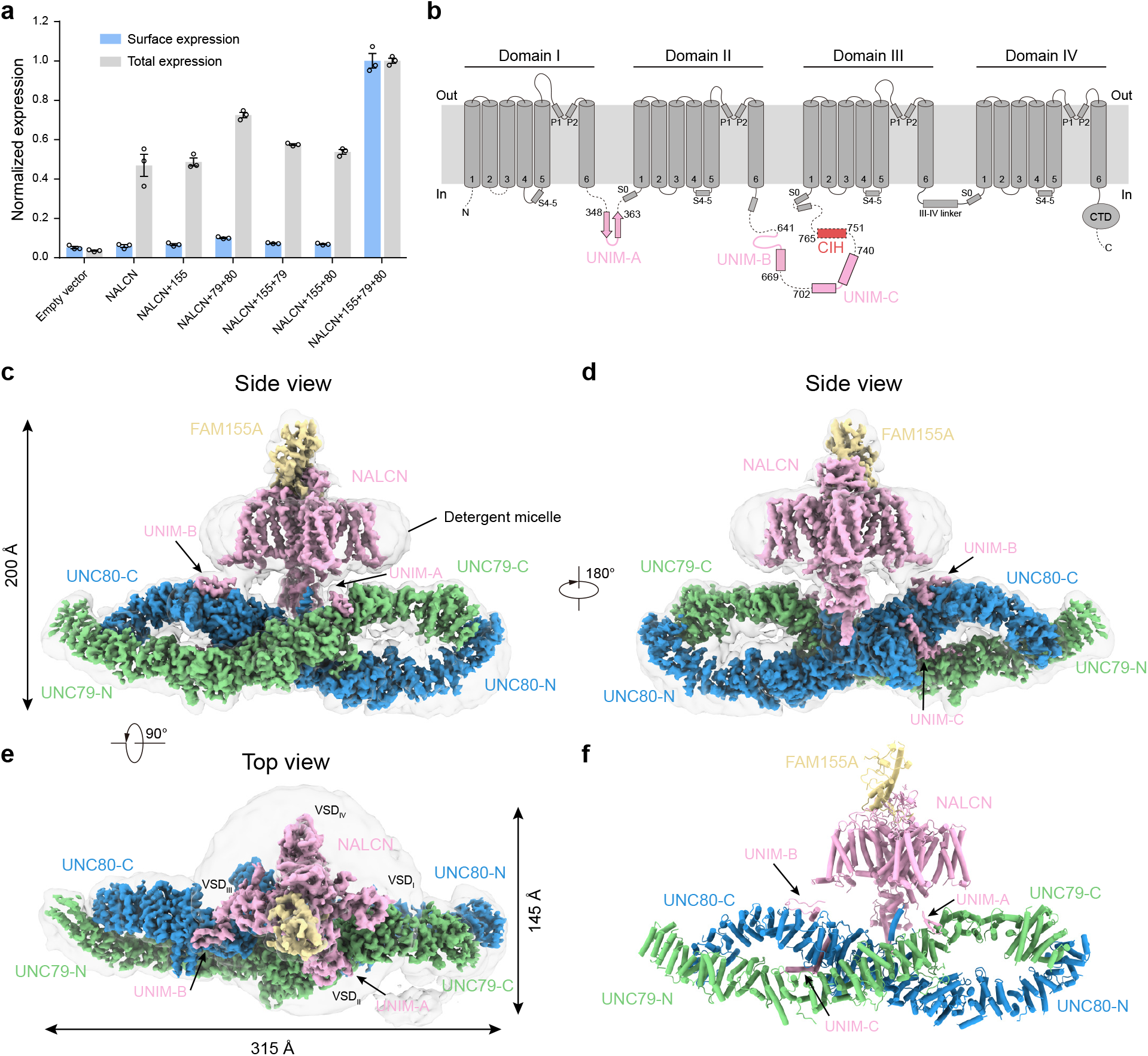
Architecture of NALCN-FAM155A-UNC79-UNC80 quaternary complex. **a** The surface localization of the NALCN subunit in the presence or absence of FAM155A, UNC79, and UNC80. The signals of the 3×HA tag on the NALCN subunit in each combination were normalized to that of co-expression of NALCN, FAM155A, UNC79, and UNC80. Data are presented as mean ± SEM, *n* = 3 biologically independent samples. **b** Topology of the NALCN subunit. The UNC-interacting motifs (UNIM) located on intracellular loops of NALCN are shown in pink. The CTD-Interacting Helix (CIH) is shown in red. **c** The side view of cryo-EM map of NALCN-FAM155A-UNC79-UNC80 quaternary complex. NALCN, FAM155A, UNC79, and UNC80 are colored in pink, yellow, green, and blue, respectively. The detergent micelle and unresolved flexible region are shown in gray and in semi-transparency. **d** A 180° rotated side view of **c**. **e** A 90° rotated top view of **c**. **f** The atomic model of NALCN-FAM155A- UNC79-UNC80 quaternary complex. Each subunit is colored the same as in **c**.

Subsequent protein purification and cryo-EM multi-body analysis showed relative motions between NALCN-FAM155A subcomplex and UNC79-UNC80 heterodimer (Supplementary information, Video S1). Further extensive focused refinements and signal subtraction resolved five overlapping structural fragments at 2.9-3.7 Å resolution (Supplementary information, Figs. S2, S3, and Table S1). The maps from focused refinement were combined to generate the composite map for model building and interpretation. We exploited AlphaFold2 to aid the manual building of UNC79 and UNC80 models ^21^. Our final model encompasses 1346 out of 1738 residues of NALCN, 173 out of 467 residues of FAM155A, 1528 out of 2654 residues of UNC79, and 1711 out of 3326 residues of UNC80 (Supplementary information, Table S1). The remaining unmodeled regions are probably highly disordered and therefore unresolvable in cryo-EM maps.

The structure of the NALCN-FAM155A subcomplex is overall similar to our previous structure at 2.65 Å (Fig. 1b-f)^15^. The UNC79-UNC80 heterodimer forms a large helix-rich structure in the cytosol. The NALCN-FAM155A subcomplex sits above the central joint of the UNC79-UNC80 heterodimer to form an asymmetric quaternary complex occupying 200 Å × 315 Å × 145 Å 3D space (Fig. 1b-f; Supplementary information, Video S2).

### Structure of the UNC79-UNC80 heterodimer

We resolved 72 cytosolic helices for UNC79 and 77 helices for UNC80 (Supplementary information, Figs. S4, S5). These cytosolic helices further pack into HAET repeats and ARM repeats which act as basic building blocks for UNC79 and UNC80 (Fig. 2a-d). We did not find any transmembrane helices of UNC79 or UNC80 in our cryo-EM density map. Although there is little sequence homology between UNC79 and UNC80, they share an overall wave-shaped structure formed by supercoiled helices (Fig. 2a-d). UNC79 and UNC80 interact in a head-to-tail fashion to form a plier-shape complex, akin to the infinity symbol “∞” (Fig. 2a-d). We divided the structures of UNC79 and UNC80 into five large consecutive functional domains: N terminal UNC-heterodimerization domain (UHD-N), UNC connecting domain 1 (UCD1), middle UNC- heterodimerization domain (UHD-M), UNC connecting domain 2 (UCD2), and C terminal UNC-heterodimerization domain (UHD-C) (Fig. 2a). UNC79 and UNC80 interact through three regions: head, joint, and tail (Fig. 2b; Supplementary information, Fig. S6). The head region is formed by 79-UHD-N and 80-UHD-C, in which α5-α13 of UNC79 interact with α71-α76 of UNC80 (Fig. 2b-d; Supplementary information, Fig. S6b). The central joint region is formed by extensive interactions between the 79-UHD-M (α26-α49) and 80-UHD-M (α24-56) with an interface of 4,270 Å^2^ area (Fig. 2b-d; Supplementary information, Fig. S6c-g). The tail region is formed by 79-UHD-C and 80-UHD-N, in which α66-α71 of UNC79 interact with α1-α8 of UNC80 (Fig. 2b-d; Supplementary information, Fig. S6h).

**Fig. 2:**
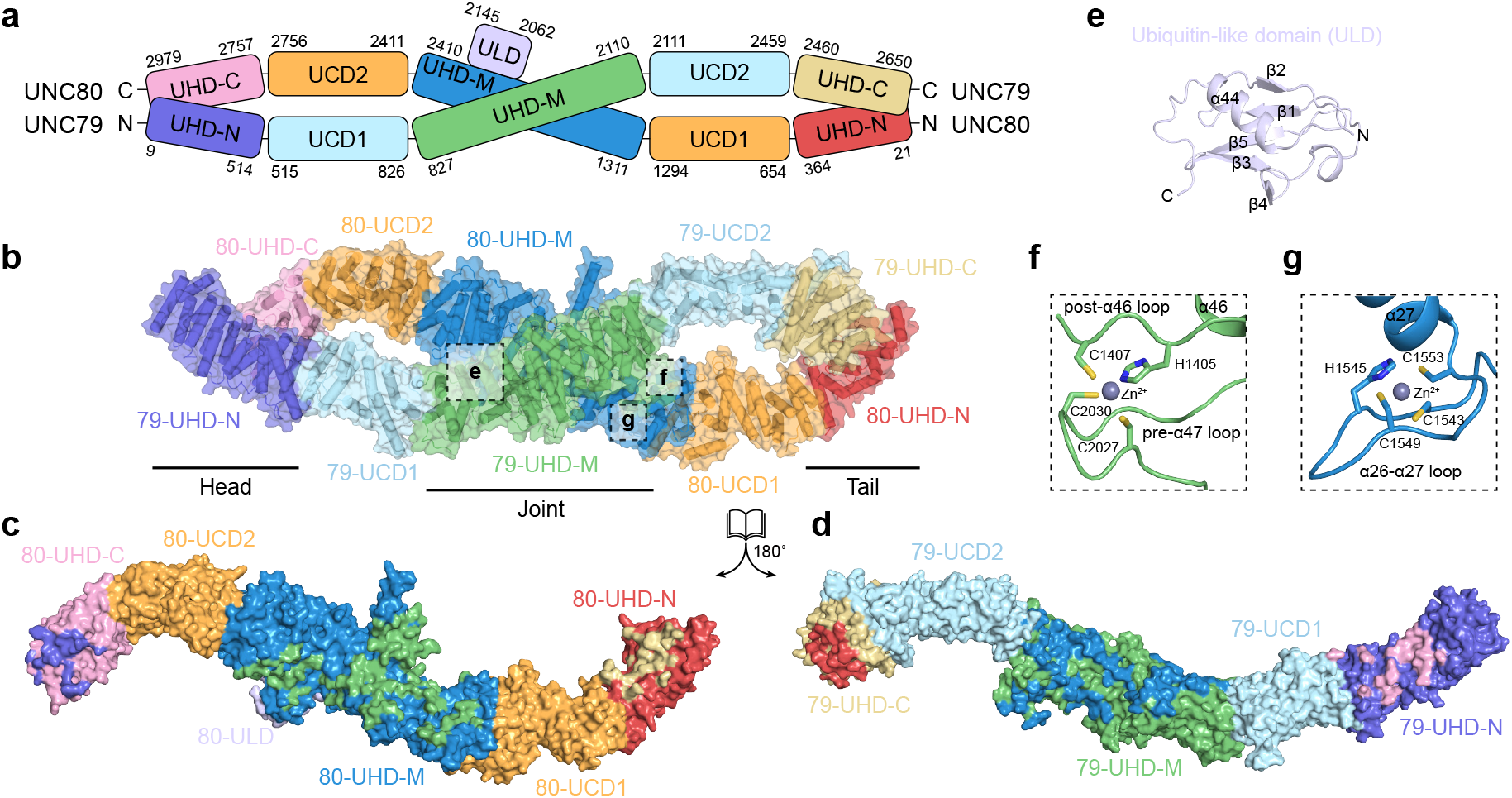
Structure of UNC79-UNC80 heterodimer. **a** Cartoon representation of the domain organizations of UNC79 and UNC80. UNC-heterodimerization domain (UHD), UNC connecting domain 1 (UCD1), middle UNC-heterodimerization domain (UHD-M), UNC connecting domain 2 (UCD2), and C terminal UNC-heterodimerization domain (UHD-C). **b** Structure of UNC79-UNC80 heterodimer colored according to domain organization shown in **a**. The helices are shown as cylinders. **c, d** The open-book view of UNC79 and UNC80 shows their interaction interface. UNC79 and UNC80 are shown in surface representation and colored according to domain organization in **a**. Interaction interfaces on each domain are stained by the colors of their interaction partners. **e** The structure of Ubiquitin-like domain (ULD) of UNC80 boxed in **b**. **f** The structure of the zinc-finger domain of UNC79 boxed in **b**. **g** The structure of the zinc-finger domain of UNC80 boxed in **b**.

Besides the HEAT and ARM repeats, there are also other structural modules present in the structures of UNC79 and UNC80. A ubiquitin-like domain (ULD) between α43 and α45 of UNC80 protrudes out of 80-UHD-M (Fig. 2e). We observed one C3H-type zinc-finger between α26 and α27 of 80-UHD-M (Fig. 2f), and another C3H-type zinc-finger between α46 and α47 of 79-UHD-M (Fig. 2g). These structural modules are located on the surface of the UNC79-UNC80 heterodimer. We further mapped the genetic mutations of UNC80 found in human patients onto the structure of rat UNC80 ^13,22–25^(Fig. 3a). We found the majority of them are involved in intra-subunit interactions of UNC80. R2901 (R2842Q in human patients) on α74 interacts with E2965 on α77 (Fig. 3b); E2634 (E2566A in human patients) interacts with K2573 on α60 (Fig. 3c); R2604 (R2536T in human patients) on α61 interacts with E2607 on α61 and E2484 close to α59 (Fig. 3d); R1724 (R1566C in human patients) on α33 interacts with E1952 on α37 (Fig. 3e). The charge-neutralization mutations found in human patients would abolish the electrostatic interactions within the UNC80 subunit and affect protein folding or overall conformation of UNC80 to impair its function, leading to diseases.

**Fig. 3:**
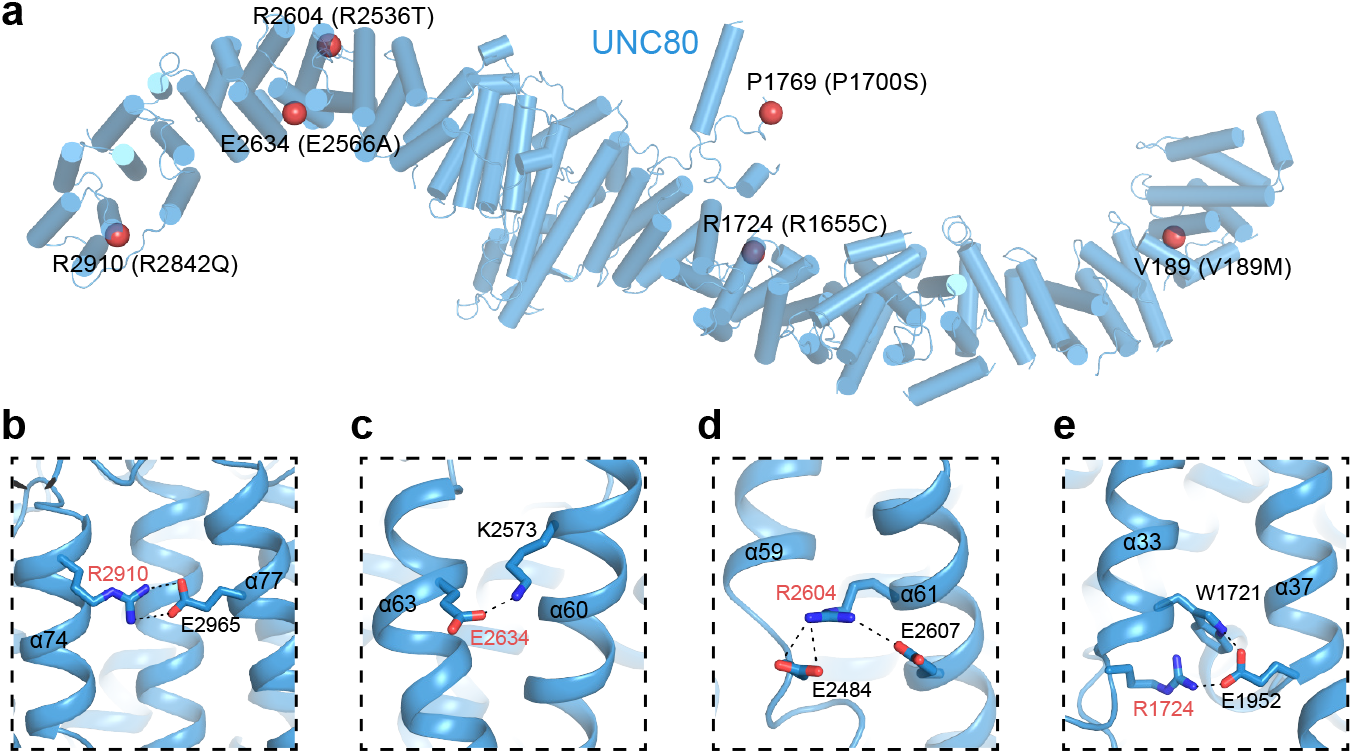
Structural mapping of disease-causing mutations in UNC80. **a** Cartoon representation of UNC80. Helices are shown as cylinders. The Cα atoms of disease-causing residues (corresponding mutations in human are indicated in parenthesis) are shown as red spheres. **b-e** Close-up view of disease-causing residue (marked with red) and its interacting residues.

### Interactions between NALCN and UNC79-UNC80 heterodimer

NALCN interacts with UNC79-UNC80 heterodimer through three cytosolic UNC-interacting motifs (UNIM-A, B, C), which all are on loop regions of NALCN (Fig. 1b). UNIM-A (residues 348-363) locates on the D_I-II_ linker of NALCN and both UNIM-B (residues 641-669) and UNIM- C (residues 702-740) reside on the D_II-III_ linker (Fig. 1b). All of these UNIMs were not resolved in previous structures of the NALCN-FAM155 subcomplex, probably due to their high flexibilities in the absence of UNC79 and UNC80. The location of UNIMs on the cytosolic loops of NALCN is consistent with the flexible tethering of NALCN-FAM155A subcomplex with the UNC79-UNC80 heterodimer, evidenced by the relative motion between them observed in the cryo-EM analysis (Supplementary information, Video S1).

UNIM-A has a short β hairpin structure which is inserted into a hydrophobic crevice formed between α48-α50 of UNC79 (Fig. 4a). In detail, F351, W359, and L361 of UNIM-A make hydrophobic interactions with A2061, L2064, L2065, and M2068 on 79-UHD-M and I2113, L2117 on 79-UCD2 (Fig. 4a). To validate the structural observation biochemically, we used co-immunoprecipitation experiments to detect the interaction between GST-tagged UNIM-A and UNC79-UNC80 heterodimer (Fig. 4b). We found UNIM-A could be co-immunoprecipitated by UNC79-UNC80 heterodimer (Fig. 4b). If the crucial hydrophobic residues on UNIM-A of NALCN (F351, W359 and L361) were mutated into alanine (UNIM-3A), the interaction was abolished (Fig. 4b).

**Fig. 4:**
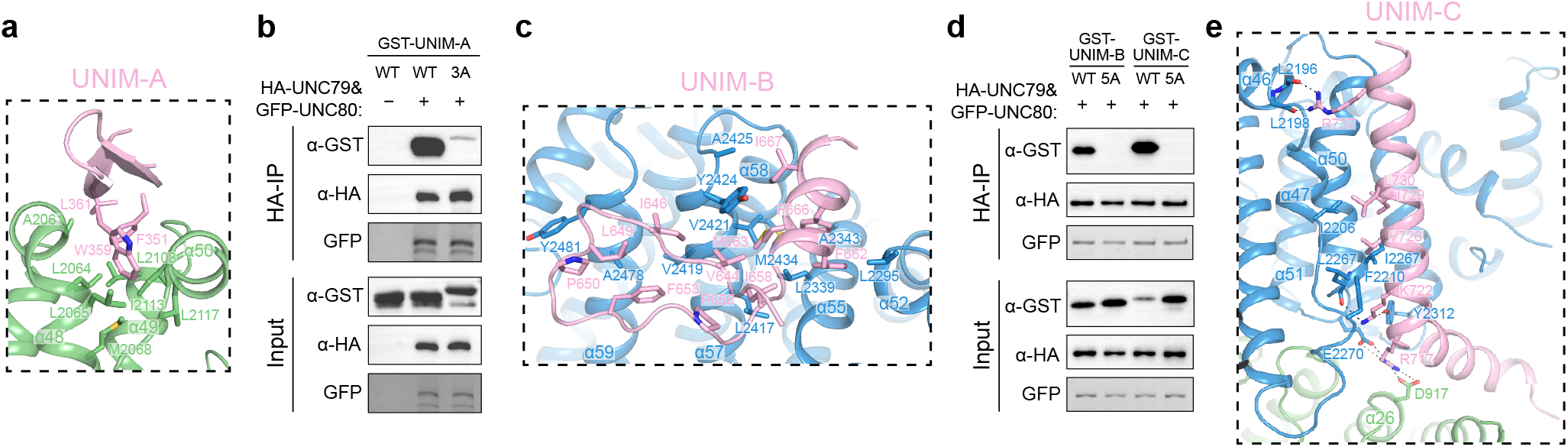
Interaction between NALCN and UNC79-UNC80 heterodimer. **a** The interaction between UNIM-A in pink and UNC79 in green. Interacting residues are shown as sticks. **b** Co-immunoprecipitation between purified HA- UNC79-GFP-UNC80 heterodimer and GST-UNIM-A or its mutants UNIM-A-3A. The experiments were repeated independently three times with similar results. **c** The interaction between UNC-interacting motif B (UNIM-B) in pink and UNC80 in blue. **d** Co-immunoprecipitation between HA-UNC79-GFP- UNC80 heterodimer and GST-UNIM-B, GST-UNIM-C, or their mutants UNIM-B-5A and UNIM-C-5A. The experiments were repeated independently three times with similar results. **e** The interaction between UNC-interacting motif C (UNIM-C) in pink and UNC80 in blue.

UNIM-B shows a loop-like structure followed by a short helix and interacts with 80-UHD-M and 80-UCD2 (Fig. 4c). Hydrophobic residues on the loop, including V644, I646, L649, P650, F653, P656, and I658, pack onto a hydrophobic patch on the surface of UNC80, making hydrophobic interactions with L2417, V2419, V2421, and Y2424 on α57, and A2478 and Y2481 on α59 (Fig. 4c). The F662, M663, F666, and I667 on the helix of UNIM-B are inserted into a hydrophobic groove formed by L2295 on α52, L2339, A2343 on α55, V2421, Y2424, A2425 on α57, and M2434 on α58 of UNC80 (Fig. 4c). Mutations of I658A, F662A, M663A, F666A, and I667A on NALCN (UNIM-B-5A) abolished the interaction between UNIM-B and UNC79-UNC80 heterodimer, evidenced by the co-immunoprecipitation experiment (Fig. 4d).

UNIM-C folds into an L-shape structure with two helices (Fig. 4e). The first helix lies on the interface between 79-UHD-M and 80-UHD-M and the second helix is sandwiched between α47 and α50 of 80-UHD-M (Fig. 4e). R717 of UNIM-C makes electrostatic interactions with D917 of 79-UHD-M and E2270 of 80-UHD-M (Fig. 4e). K722 of UNIM-C makes polar interactions with Y2312 and main-chain carbonyl group of L2267 of 80-UHD-M (Fig. 4e). V726, I729, and L730 of UNIM-C make hydrophobic interactions with I2206, F2210, I2266, and L2267 of 80- UHD-M (Fig. 4e). R737 makes hydrogen bonding with the main-chain carbonyl group of L2196 and L2198 of 80-UHD-M (Fig. 4e). Mutations of R717A, K722A, V726A, I729A, and L730A on NALCN (UNIM-C-5A) diminished the binding of UNIM-C to UNC79-UNC80 (Fig. 4d).

### Mechanism of NALCN channel regulation by UNC79-UNC80

The UNC79-UNC80 heterodimer regulates the NALCN channel through physical interactions with UNIMs of NALCN observed in the structure. In order to study the functional roles of these three identified interactions, we disrupted them individually by making the aforementioned mutations on UNIMs and recorded the NALCN currents of the mutants. We found mutations of UNIM-A (UNIM-A-3A) and UNIM-B (UNIM-B-5A) diminished the whole-cell currents of NALCN (Fig. 5a, b). The reduction of currents correlates with the reduced surface localization of NALCN (Fig. 5c). Therefore, we speculate that there are cytoplasmic retention signals, such as ER retention signals, on cytosolic regions of NALCN, and the binding of UNIM-A and UNIM-B onto UNC79-UNC80 heterodimer masks such signals to facilitate the surface localization of the NALCN channel. This is akin to the K_ATP_ channel, in which the SUR subunits mask the ER retention signal of Kir6 subunits to promote the surface localization of the fully assembled K_ATP_ channel ^26^.

**Fig. 5:**
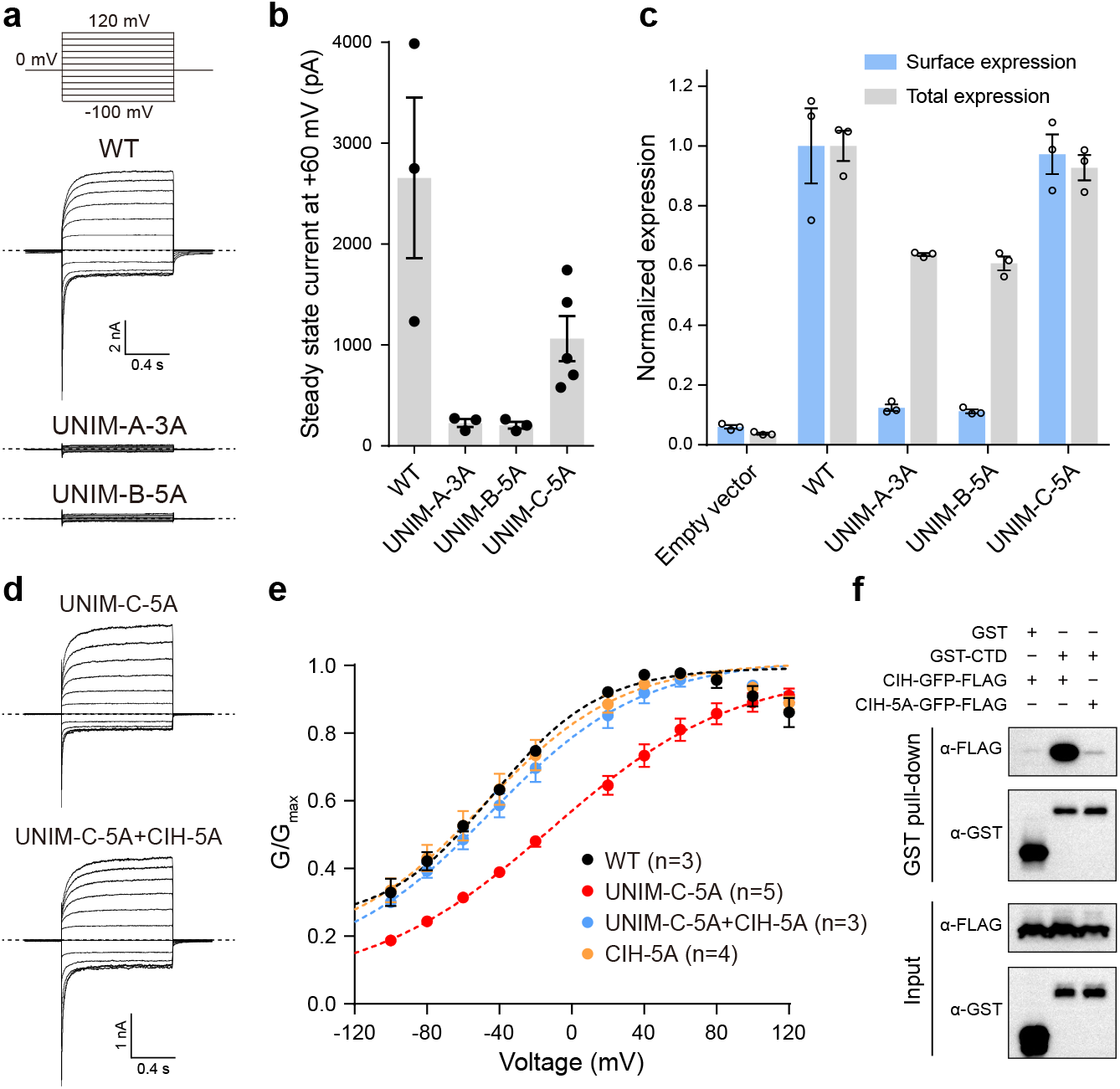
Mechanism of NALCN activation by UNC79-UNC80 heterodimer. **a** The whole-cell voltage step protocol and representative whole-cell current traces of wild-type NALCN, UNIM-A-3A, or UNIM-B-5A mutants in the presence of FAM155A, UNC79, and UNC80. Dashed lines indicate the position of 0 nA. The experiments were repeated independently three times with similar results. **b** Whole-cell steady-state currents of wild-type NALCN, UNIM-A-3A, UNIM-B-5A, or UNIM-C-5A mutants in the presence of FAM155A, UNC79, and UNC80 at +60 mV. Data are presented as mean ± SEM, *n* ≥ 3 biologically independent cells. **c** The normalized surface localization of the NALCN subunit and its mutants in the presence of FAM155A, UNC79, and UNC80. Data are presented as mean ± SEM, *n* = 3 biologically independent samples. **d** Representative whole-cell current traces of UNIM-C-5A and UNIM-C-5A+CIH-5A mutants of NALCN in the presence of FAM155A, UNC79, and UNC80. The whole-cell voltage step protocol was same as in **a**. Dashed lines indicate the position of 0 nA. The experiments were repeated independently at least three times with similar results. **e** Conductance versus voltage (G-V) curve of wild-type NALCN and its mutants in the presence of FAM155A, UNC79, and UNC80. The conductance at each voltage step was normalized to the maximum conductance obtained by fitting the data into the Boltzmann equation. Data are presented as mean ± SEM, *n* ≥ 3 biologically independent cells. **f** The GST pull-down assay of interactions between NALCN-CTD and CIH-GFP or its mutant CIH-5A. The experiments were repeated independently three times with similar results.

In contrast, the UNIM-C-5A mutant retains both robust whole-cell currents and surface localization compared to the wild-type channel (Fig. 5c, d). However, the conductance versus voltage (G-V) curve of the UNIM-C-5A mutant shifts to the positive potential compared with the wild-type channel (Fig. 5e), suggesting that the UNIM-C-5A mutation inhibits the opening of the NALCN. Close inspection of the cryo-EM map showed that the CIH density was missing in the NALCN-FAM155A-UNC79-UNC80 quaternary complex (Supplementary information, Fig. S7a). Both UNIM-C (702-740) and CIH (751-765) are on the D_II-III_ linker of NALCN and there are only 10 amino acids between UNIM-C and CIH. If we modeled the CIH in its binding site of NALCN CTD of the quaternary complex, we found the linear distance between the termini of UNIM-C and CIH is about 60 Å (Supplementary information, Fig. S7b), which is much longer than the distance that 10 amino acids could extend to, even in their fully extended conformation. Therefore, our structural observation and modeling study highly suggest the possibility that the binding of UNIM-C of NALCN D_II-III_ linker to UNC79-UNC80 heterodimer in the quaternary complex would pull the neighboring CIH out of the binding pocket in NALCN CTD, and UMIM-C-5A mutations would disrupt the binding of UNIM-C to UNC79-UNC80 and release the pulling force on CIH to allow it to re-bind to NALCN CTD. The rightward shift of the G-V curve of the UNIM-C-5A mutant indicates that the binding of CIH in CTD inhibits channel opening, probably by stabilizing the channel in a closed state. To further validate this model, we looked for the mutants on CIH which could disrupt the interaction between CIH and CTD. Guided by our previous high resolution structure of NALCN-FAM155A subcomplex ^15^, we found that the combination of I753A, L754A, R761A, R764A and R765A mutations (CIH-5A) is sufficient to disrupt such interaction, shown by GST pull-down assay (Fig. 5f). We hypothesis that the CIH-5A mutation might release the self-inhibition of NALCN. Indeed, we found that additional CIH-5A mutation on the background of UNIM-C-5A mutant shifts the G-V curve of UNIM-C-5A back to the level similar to the wild-type channel, while CIH-5A itself has no effect on the gating of NALCN (Fig. 5e), in agreement with our mechanistic model.

In summary, the near-atomic resolution structure of the NALCN-FAM155A-UNC79-UNC80 complex presented here shows the architecture of the functional mammalian NALCN channel complex, reveals the structure of UNC79-UNC80 heterodimer, and depicts the detailed interactions between UNC79 and UNC80. More importantly, the structure uncovers the activation mechanism of the NALCN channel by UNC79-UNC80 heterodimer, not only through promoting the surface localization of NALCN but also through modulating the gating of NALCN (Fig. 6). Our work provides a structural basis to target the NALCN channel complex for future therapeutic intervention.

**Fig. 6:**
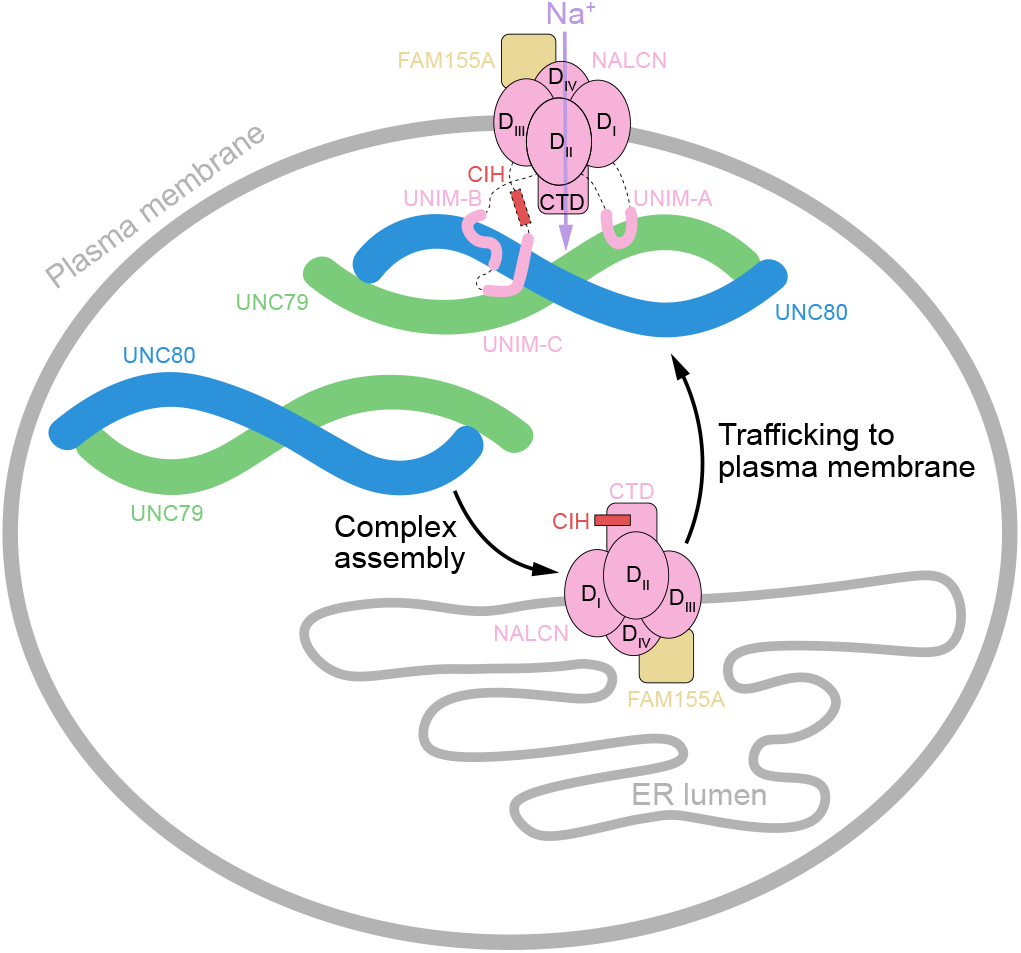
Mechanistic model of NALCN-FAM155A-UNC79-UNC80 channel complex. NALCN, FAM155A, UNC79, and UNC80 are shown as cartoons and colored the same as Fig. 1c. Initially, the NALCN-FAM155A subcomplex is assembled in an intracellular compartment, probably in the ER. Subsequently, the UNC79-UNC80 heterodimer binds to the UNIM-A, B, and C of NALCN-FAM155A subcomplex to activate the channel. Binding of UNC79-UNC80 with UNIM-A and UNIM-B promotes plasma localization of NALCN complex. Binding of UNC79-UNC80 with UNIM-C releases the auto-inhibition of NALCN by CIH.

---

**Video S1. The relative motion between NALCN-FAM155A subcomplex and UNC79- UNC80 heterodimer revealed by multi-body analysis.**

Repositioning of reconstructed densities of NALCN-FAM155A and UNC79-UNC80 along the first and the second eigenvectors reveals the relative motions between NALCN-FAM155A and UNC79-UNC80. Eigenvector #1 explains 29% variance and eigenvector #2 explains 25% variance.

**Video S2. Overall structure of NALCN-FAM155A-UNC79-UNC80 quaternary complex.**

The cryo-EM map of NALCN-FAM155A-UNC79-UNC80 quaternary complex.

## Methods

### Cell Culture

Sf9 insect cells (Thermo Fisher Scientific, Waltham, MA, USA) were cultured in SIM SF (Sino Biological, China) at 27 °C. FreeStyle 293F suspension cells (Thermo Fisher Scientific) were cultured in FreeStyle 293 medium (Thermo Fisher Scientific) supplemented with 1% fetal bovine serum (FBS) at 37 °C with 6% CO_2_ and 70% humidity. HEK293T (ATCC) were cultured in Dulbecco’s Modified Eagle Medium (Gibco, Gaithersburg, MD USA) supplemented with 10% FBS at 37 °C with 5% CO_2_. The cell lines were routinely checked to be negative for mycoplasma contamination but have not been authenticated.

### Determination of the surface expression of NALCN subunit

HEK293T cells were plated onto poly-D-lysine treated 24-well plate (303002; Porvair Sciences) and transfected with indicated plasmids (Figs. 1a, 5c, in a modified BacMam expression vector ^27^) at a ratio of 1:1.5:1.75:2 (FAM155A-3×FLAG:NALCN-1077-3×HA-GFP:UNC79-3×FLAG:UNC80-3×FLAG) using Lipofectamine 3000 (Thermo Fisher Scientific) and incubated for 40-48 hours. When certain constructs were excluded from the transfection, an equal amount of empty vector was added. The cells were washed with PBS twice, fixed using 4% formaldehyde in PBS for 30 min, and washed with PBS twice again. Then the cells were blocked with 3% goat serum in PBS for 30 min and labeled with primary antibody [rabbit anti-HA (3724; CST, diluted 2,500 times in blocking buffer)] for 1 hour. After washing with PBS for 3 times, cells were incubated with horseradish-peroxidase (HRP) labeled goat anti-rabbit secondary antibody (31460; Thermo Fisher Scientific, the antibody was diluted 2,500 times in blocking buffer) for 30 min. After extensive washing, the cells were incubated with High-Sig ECL Western Blotting Substrate (Tanon) for 2 min, and chemiluminescence signals were measured with Infinite M Plex plate reader (Tecan). The signals represent the surface expression of the NALCN subunit. Then the cells were permeabilized by incubating in PBS with 0.1% TX-100 for 30 min. The cells were subsequently blocked, labelled with primary and secondary antibodies, and detected by ECL chemiluminescence as described above. The chemiluminescence signals of permeabilized cells represents the total expression of NALCN.

### Fluorescence-detection size-exclusion chromatography (FSEC)

Purified protein or cell lysates were injected onto a Superose 6 increase 5/150 column (GE Healthcare), running in a buffer containing 20 mM Tris (pH 7.5), 150 mM NaCl, 0.5 mM *n*-dodecyl β-D-maltoside (DDM, Anatrace), and detected by a fluorescence detector (Shimadzu, excitation 488 nm and emission 520 nm for GFP signal) at room temperature.

### Electrophysiology

HEK293T cells were co-transfected with constructs containing rNALCN-GFP, mFAM155A- FLAG, mScarlet-mUNC80, and HA-mUNC79 plasmids or their various mutants (in a modified BacMam expression vector ^27^) at a ratio of 2:1:1:1 using Lipofectamine 3000 (Thermo Fisher Scientific) and incubated for 24 hours before recording. Patch electrodes were pulled with a horizontal microelectrode puller (P-1000, Sutter Instrument Co, USA) to a resistance of 1.7-2.5 MΩ. Whole-cell patch clamps were performed using an Axon-patch 200B amplifier (Axon Instruments, USA), and data were collected with pClamp 10 software (Axon Instruments, USA) and an Axon Digidata 1550B digitizer (Axon Instruments, USA). Pipette solution containing (mM): 10 HEPES (pH 7.2, NaOH), 136 NaCl, 10 NaF, 5 EGTA, 2 Na_2_ATP and bath solution containing (mM): 10 HEPES (pH 7.4, NaOH), 150 NaCl, 30 glucose. At the end of each experiment, the bath solution was exchanged to 10 mM HEPES (pH 7.4, HCl), 150 mM NMDG, and 30 mM glucose by local perfusion system (MPS-2, InBio) to exclude the loosely sealed patch (cells with current <−100 pA at −100 mV was discarded). Serial resistance was compensated by at least 75%. The steady-state currents were used to generate G/G_max_ versus V curves (G□=□I/V). The Boltzmann equation was used to fit the G-V curve in GraphPad Prism 6. Signals were acquired at 5 kHz and low-pass filtered at 1 kHz. The data was processed using Clampfit, Microsoft Excel and GraphPad Prism 6.

### Protein expression and purification

The cDNAs of rat NALCN, mouse FAM155A, mouse UNC79, and mouse UNC80 were cloned into a modified BacMam expression vector ^27^ with C-terminal GFP-strep-FLAG tag, a FLAG tag, N-terminal HA tag and NbGFP-strep-FLAG tag, respectively. The baculoviruses were produced using the Bac-to-Bac system and amplified in Sf9 cells. We expressed NALCN-FAM155A subcomplex and UNC79-UNC80 heterodimer separately. For protein expression, FreeStyle 293F cells cultured in FreeStyle 293 medium at a density of 2.5 × 10^6^ ml^-1^ were infected with 6% volume of NALCN P2 virus and 4% volume of FAM155A P2 virus or 6% volume of UNC79 P2 virus and 4% volume of UNC80 P2 virus. 10 mM sodium butyrate was added to the culture 12 hours post-infection and transferred to a 30°C incubator for another 48 hours (NALCN- FAM155A) or 60 hours (UNC79-UNC80) before harvesting. Cells were collected by centrifugation at 4,000 rpm (JLA-8.1000, Beckman) for 10 min, and washed with 20 mM Tris (pH 7.5), 150 mM NaCl, 2 μg/ml aprotinin, 2 μg/ml pepstatin, and 2 μg/ml leupeptin, flash frozen and storage at −80°C.

For each batch of protein purification, cell pellets corresponding to 0.6 liter culture of NALCN- FAM155A or UNC79-UNC80 was thawed and extracted with 42 ml or 21 ml lysis buffer containing 50 mM HEPES (pH 8.0), 150 mM NaCl, 2 μg/ml aprotinin, 2 μg/ml pepstatin, 2 μg/ml leupeptin, 10% (v/v) glycerol, 1 mM phenylmethanesulfonyl fluoride (PMSF), 2 mM MgCl_2_, 0.7 μg/ml benzonase, 1mg/ml iodoacetamide and 1% (w/v) glyco-diosgenin (GDN, Anatrace) at 4°C for 1 hour, respectively. The lysates were then mixed and incubated for 30 min and centrifuged at 40,000 g (JA25.5, Beckman) for 40 min at 4°C. The supernatant was incubated with 1 ml Anti-FLAG Affinity Beads (Smart-Lifesciences) for 90 min at 4°C and washed with 5 ml W buffer (20 mM HEPES (pH 8.0), 150 mM NaCl, 10% glycerol, 2 μg/ml aprotinin, 2 μg/ml pepstatin, 2 μg/ml leupeptin, and 0.02% GDN) for 4 times. The target protein was eluted with 1ml W buffer plus 200 μg/ml 3×FLAG peptides (Smart-Lifesciences) and 30 mM HEPES (pH 8.0) 5 times. The eluate was loaded onto 2ml Streptactin Beads 4FF (Smart-Lifesciences) column and washed with 45 ml W buffer plus 2 mM ATP and 10 mM MgCl_2_, and washed with 20 mM HEPES (pH 8.0), 150 mM NaCl. 10% glycerol and 0.02% GDN for 20 ml. The target protein was eluted by 25 mM HEPES (pH 8.5), 150 mM NaCl, 10% glycerol, 0.02% GDN and 5 mM desthiobiotin (IBA). Eluted protein was crosslinked with 0.04% glutaraldehyde (EM-grade, Sigma) on ice for 25 min, and stopped with 50 mM Tris-HCl (pH 7.5) on ice for 15 min. Crosslinked protein was concentrated using 100-kDa cut-off concentrator (Millipore) and further purified by Superose 6 increase (GE Healthcare) running in a buffer containing 20 mM Tris (pH 7.5), 150 mM NaCl and 0.006% GDN. Fractions corresponding to NALCN-FAM155A-UNC79- UNC80 complex were pooled and concentrated to A_280_ = 0.38 for cryo-EM sample preparation.

### Cryo-EM sample preparation and data collection

Holey carbon grids (Quantifoil Au 300 mesh, R 0.6/1) coated with home-made ultrathin continuous carbon were glow-discharged by plasma cleaner (Harrick, PDC-32G) for 30 s with the “Low” setting. Aliquots of 3 μl concentrated protein sample were applied on glow-discharged grids and the grids were blotted for 2 s before plunged into liquid ethane using Vitrobot Mark IV (Thermo Fisher Scientific). Cryo-grids were firstly screened on a Talos Arctica electron microscope (Thermo Fisher Scientific) operating at 200 kV with a K2 Summit direct electron camera (Thermo Fisher Scientific). The screened grids were subsequently transferred to a Titan Krios electron microscope (Thermo Fisher Scientific) operating at 300 kV with a K2 Summit direct electron camera and a GIF Quantum energy filter set to a slit width of 20 eV. Images were automatically collected using SerialEM in super-resolution mode at a nominal magnification of 105,000 ×, corresponding to a calibrated super-resolution pixel size of 0.662 Å with a preset defocus range from −1.5 μm to −1.8 μm. Each image was acquired as a 10 s movie stack of 40 frames with a dose rate of 5 e^-^Å^−2^s^−1^, resulting in a total dose of about 50 e^−^Å^−2^.

### Cryo-EM image analysis

The image processing workflow is illustrated in Supplementary information, Fig. S2c. A total of 6,714 super-resolution movie stacks were collected. Motion-correction, two-fold binning to a pixel size of 1.324 Å, and dose weighting were performed using MotionCor2 ^28^. Contrast transfer function (CTF) parameters were estimated with Gctf ^29^. Micrographs with ice or ethane contamination and empty carbon were removed manually. A total of 2,742,149 particles were auto-picked using Gautomatch from 6,156 micrographs. All subsequent classification and reconstruction were performed in Relion 3.1 ^30^ unless otherwise stated. Two rounds of reference- free 2D classification were performed to remove contaminants, yielding 797,032 particles. The particles were subjected to 30 iterations K = 1 global search 3D classification with an angular sampling step of 7.5° to determine the initial alignment parameters using the initial model generated from cryoSPARC ^31^. For each of the last five iterations of the global search, a K = 4 multi-reference (resolution gradient) local angular search 3D classification with the mask of UNC79-UNC80 was performed with an angular sampling step of 3.75° and search range of 30°. The classes that showed obvious secondary structure features were selected and combined. Duplicated particles were removed, yielding 567,778 particles in total. These particles were subjected to additional six rounds of multi-reference local angular search 3D classification with the mask of UNC79-UNC80 using resolution gradient references or good and phase randomized references, yielding 202,781 particles. These particles were subsequently subjected to local NU- refinement in cryoSPARC with the mask of UNC79-UNC80 ^32^, which resulted in a map with a resolution of 3.3 Å. To further improve the resolution, seed-facilitated 3D classification was performed ^33^. The CTF parameters were re-estimated with Patch CTF in cryoSPARC. 486,791 particles were selected after two rounds of seed-facilitated 3D classification using good and biased references or references with resolution gradients in cryoSPARC. These particles were subjected to reference-free 2D classification, yielding 430,068 particles. The particles were subjected to global and local search 3D classification to determine the correct alignment parameters, yielding 336,476 particles. In further clean up the dataset, no alignment 3D classification was performed with the mask of the UNC79-UNC80 middle region. Particles with the best features were selected, yielding 275,170 particles. These particles were subjected to local NU-refinement in cryoSPARC with the mask of UNC79- UNC80 or whole complex, which resulted in maps with resolutions of 3.1 Å and 3.2 Å, respectively. We applied different local masks of UNC79-UNC80 fragments to improve the local map quality, yielding four overlapping focus-refined maps with resolutions of 2.9-3.4 Å. Due to the relative motions between NALCN- FAM155A and UNC79-UNC80, we performed particle subtraction to remove signals of UNC79- UNC80 to improve the alignment of the NALCN-FAM155A subcomplex. After 3 rounds of local search 3D classification using subtracted particles, 68,126 particles were selected. These particles were subjected to local NU-refinement in cryoSPARC with the mask of NALCN- FAM155A, yielding a map with a resolution of 3.7 Å. These subtracted particles were re-extracted on original micrographs and analyzed by multi-body refinement with masks of NALCN-FAM155A and UNC79-UNC80 ^20^. The focus-refined maps were combined to generate the composite map for model building and interpretation.

All of the resolution estimations were based on a Fourier shell correlation (FSC) of 0.143 cutoff after correction of the masking effect. B-factor used for map sharpening was automatically determined by NU-refinement in cryoSPARC. The local resolution was estimated with Relion 3.1 with half maps output from cryoSPARC.

### Model building

The model of NALCN-FAM155A (PDB ID: 7CU3) was docked into the electron density map using UCSF Chimera ^34^ and manually adjusted using Coot ^35^. The models of the UNC79-UNC80 joint region were built manually using Coot. The models of UNC79-UNC80 head and tail regions were predicted by AlphaFold2 ^21^, docked into cryo-EM map and manually adjusted using Coot. The initial models of UNIMs were traced by DeepTracer ^36^ according to the cryo-EM maps and protein sequences, and re-built manually using Coot. Model refinement was performed using phenix.real_space_refine in PHENIX ^37^. Images were produced using Pymol, UCSF Chimera^34^ and ChimeraX^38^. Sequence alignment was performed with ClustalX2^39^ and illustrated by ESPript 3.0 ^40^.

### Co-immunoprecipitation

For co-immunoprecipitation between UNIM-A and UNC79-UNC80 (Fig. 4b), fragment of UNIM-A WT or 3A mutant was cloned into a pGEX-6P-1 vector. GST-UNIM-A WT or 3A was over-expressed in *E. coli* NiCo21 (DE3) induced with 1 mM isopropyl-β-D-thiogalactoside (IPTG) when the cell density reached OD_600_ = 0.6 for 4 hours at 37°C. The cells were collected and sonicated in lysis buffer containing 50 mM Tris (pH 7.5), 150 mM NaCl, 2 μg/ml aprotinin, 2 μg/ml pepstatin, 2 μg/ml leupeptin, and 1 mM PMSF. Then the cell debris was removed by centrifugation at 30,966×g (JA-25.50, Beckman) for 30 min. the supernatants were loaded onto Glutathione Sepharose 4B and washed with 20 mM Tris (pH 7.5), 150 mM NaCl and eluted with 50 mM Tris (pH 8.0), 50 mM NaCl and 10 mM reduced glutathione. GST eluates were further purified with an anion exchange column (HiTrap Q HP, GE Healthcare) and the peak fractions were pooled for co-immunoprecipitation experiments. To obtain the protein of UNC79-UNC80 heterodimer, we expressed and purified N-terminal HA tagged UNC79 and GFP-strep tagged UNC80 by infecting FreeStyle 293F cells with corresponding BacMam viruses. Protein was purified with Streptactin affinity beads as described in ‘Protein expression and purification’ section. 200 μl of purified GST-UNIM-A-WT/3A (A_280_ = 0.744, diluted by 20 mM Tris (pH 7.5), 150 mM NaCl, 10% glycerol, 2 μg/ml aprotinin, 2 μg/ml pepstatin, 2 μg/ml leupeptin and 0.006% GDN) was mixed with 100 μl purified HA-UNC79 and GFP-UNC80 heterodimer (A_280_ = 0.138 in Streptactin elution buffer, 20 μl of the mixture was saved as input) and incubated with Anti-HA Magnetic Beads (88836; Thermo Fisher Scientific) for 2 hours at 4°C. The beads were washed with 20 mM Tris (pH 7.5), 80 mM NaCl, 2 μg/ml aprotinin, 2 μg/ml pepstatin, 2 μg/ml leupeptin, 10% glycerol and 0.006%GDN 5 times, and eluted with 30 μl SDS-PAGE loading buffer (without reducing agent).

For co-immunoprecipitation of UNIM-B/C and UNC79-UNC80 (Fig. 4d), cDNA of UNIM-B/C WT or 5A mutant was cloned into a modified BacMam expression vector ^27^. FreeStyle 293F cells cultured in FreeStyle 293 medium were transfected with indicated plasmids (Fig. 4d) at a ratio of 1:1:1 using PEI. 48 hours post-transfection, 2 ml cells were collected and lysed using 300 μl 20 mM Tris (pH 7.5), 150 mM NaCl, 2 μg/ml aprotinin, 2 μg/ml pepstatin, 2 μg/ml leupeptin, 1 mM PMSF, 20% glycerol and 0.5% GDN at 4 °C for 30 min. After ultra-centrifugation at 100,000×g (TLA-55, Beckman) for 30 min, the supernatants (20 μl of lysate was saved as input) were mixed with Anti-HA Magnetic Beads (88836; Thermo Fisher Scientific) and incubated at 4 °C for 2.5 hours. Then the beads were washed with 20 mM Tris (pH 7.5), 150 mM NaCl, 2 μg/ml aprotinin, 2 μg/ml pepstatin, 2 μg/ml leupeptin, 20% glycerol and 0.006% GDN 5 times. Bound proteins were eluted with 30 μl SDS-PAGE loading buffer (without reducing agent).

Proteins of input and elution were separated with SDS-PAGE and transferred onto polyvinylidene difluoride (PVDF) membranes (GFP signal was detected by in-gel fluorescence). PVDF membranes were blocked using 5% nonfat milk in TBST [25mM Tris (pH 7.4), 137 mM NaCl, 3 mM KCl and 0.1% Tween-20] for 1 hour at room temperature and incubated with primary antibodies [mouse anti-GST (30901ES10; Yeasen Biotechnology) or [rabbit anti-HA (3724; CST), both of antibodies were diluted 5,000 times] overnight at 4°C. Then membranes were incubated with corresponding horseradish-peroxidase (HRP) labeled goat anti-mouse secondary antibody (31444; Thermo Fisher Scientific) or horseradish-peroxidase (HRP) labeled goat anti-rabbit secondary antibody (31460; Thermo Fisher Scientific, both of antibodies were diluted 10,000 times) for 1 hour at room temperature and developed using High-Sig ECL Western Blotting Substrate (Tanon).

### GST pull-down assay

FreeStyle 293F cells cultured in FreeStyle 293 medium were transfected with indicated plasmids (Fig. 5f) at a ratio of 1:1 using PEI. 48 hours post-transfection, cells were collected and sonicated in a buffer containing 20 mM Tris (pH 7.5), 150 mM NaCl, 2 μg/ml aprotinin, 2 μg/ml pepstatin, 2 μg/ml leupeptin, 1 mM PMSF. After centrifugation at 14,800 rpm for 20 min, the supernatants were mixed with Glutathione Sepharose 4B (GE Healthcare) and incubated at 4°C. for 1 hour. Then the beads were washed with 20 mM Tris (pH 7.5) and 150 mM NaCl 6 times. Bound proteins were eluted with 50 mM Tris (pH 8.0), 150 mM NaCl, and 10 mM reduced glutathione.

For Western blot, proteins were separated with 10% SDS-PAGE and transferred onto PVDF membranes. Membranes were blocked using 5% nonfat milk in TBST for 1 hour at room temperature and incubated with primary antibodies [mouse anti-GST (30901ES10; Yeasen Biotechnology) or mouse anti-FLAG (M20008M; Abmart), both of antibodies were diluted 5,000 times] overnight at 4°C. Then membranes were incubated with horseradish-peroxidase (HRP) labeled goat anti-mouse secondary antibody (31444; Thermo Fisher Scientific, the antibody was diluted 10,000 times) for 1 hour at room temperature and developed using High-Sig ECL Western Blotting Substrate (Tanon).

### Quantification and statistical analysis

Global resolution estimations of cryo-EM density maps are based on the 0.143 Fourier Shell Correlation criterion ^41^. The local resolution was estimated using Relion3.1 with half maps output from cryoSPARC. The number of independent experiments (N) and the relevant statistical parameters for each experiment (such as mean or standard error of mean) are described in the figure legends. No statistical methods were used to pre-determine sample sizes.

## Acknowledgements

We thank Prof. Dejian Ren for providing the cDNA of hNALCN, rNALCN, mUNC80, and mUNC79. Cryo-EM data collection was supported by Electron microscopy laboratory and Cryo-EM platform of Peking University with the assistance of Xuemei Li, Zhenxi Guo, Bo Shao, Xia Pei and Guopeng Wang. Part of the structural computation was also performed on the Computing Platform of the Center for Life Science and High-performance Computing Platform of Peking University. We thank the National Center for Protein Sciences at Peking University in Beijing, China for assistance with negative stain EM. The work is supported by grants from the National Natural Science Foundation of China (91957201, 31870833, and 31821091 to L.C.). Y.K. is supported by the Boya Postdoctoral Fellowship of Peking University.

## Author contributions

L.C. initiated the project and wrote the manuscript draft. Y.K. carried out experiments with the help of L.C.. Both authors contributed to the manuscript preparation.

## Data availability

Cryo-EM maps and atomic coordinates of the NALCN-FAM155A-UNC79-UNC80 complex have been deposited in the EMDB and PDB under the ID codes EMDB: EMD-32344 and PDB: 7W7G, respectively.

## Additional Information

**Correspondence and requests for materials** should be addressed to L.C.

---

**Fig. S1:**
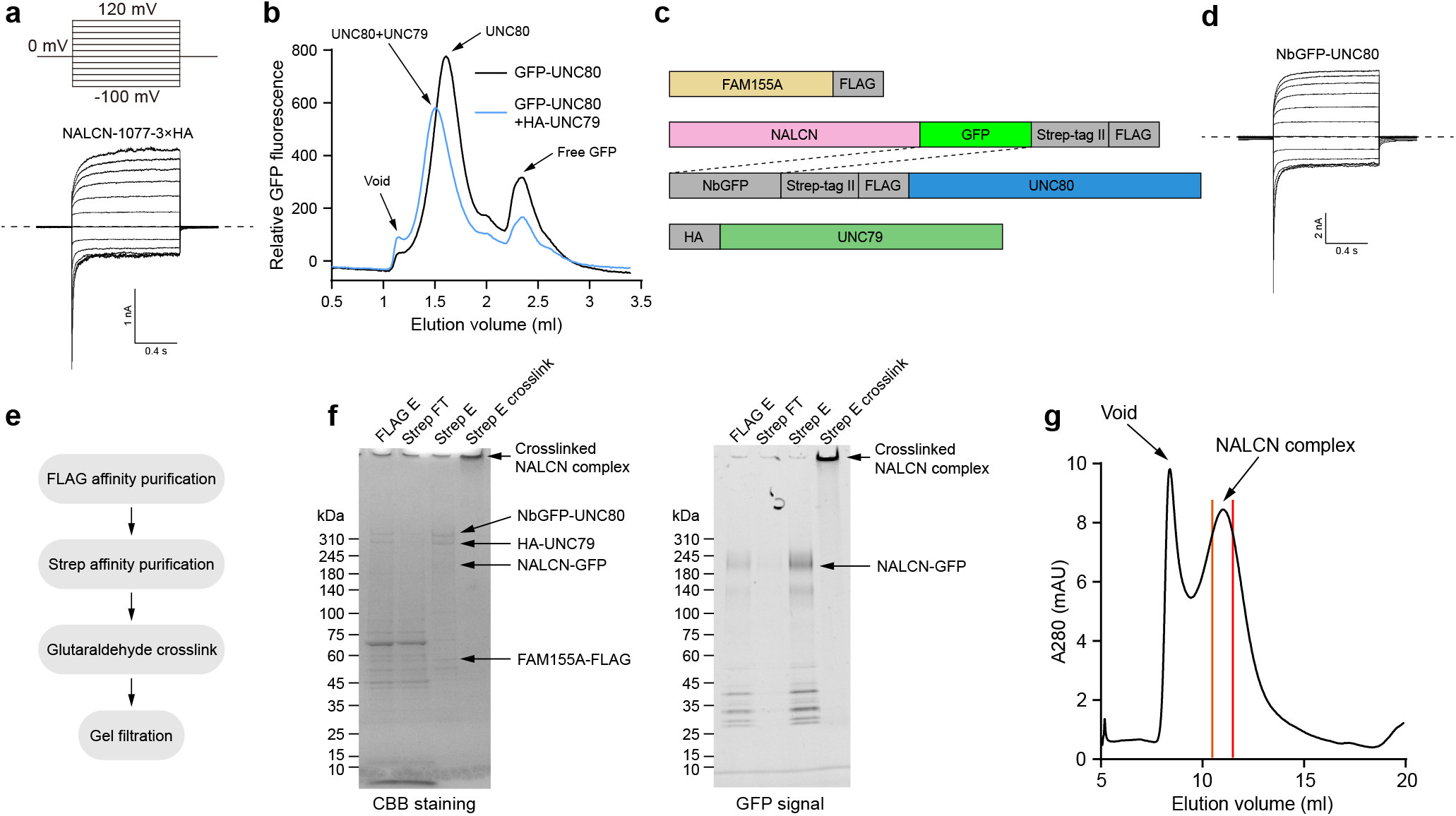
Electrophysiology and biochemistry characterization of NALCN-FAM155A-UNC79-UNC80 quaternary complex. **a** The whole-cell voltage step protocol and representative whole-cell currents of NALCN with 3×HA tag inserted between residues 1077 and 1078 in the presence of FAM155A, UNC79, and UNC80. Dashed lines indicate the position of 0 nA. The experiments were repeated independently three times with similar results. **b** Fluorescence-detection size-exclusion chromatography of GFP-UNC80 in the absence (black) or presence of UNC79 (blue). The experiments were repeated independently three times with similar results. **c** The construct used for protein purification. **d** Representative whole-cell currents of constructs shown in **c**. The whole-cell voltage step protocol was same as in **a**. Dashed lines indicate the position of 0 nA. The experiments were repeated independently three times with similar results. **e** The protein purification work-flow. **f** SDS-PAGE of protein purified by affinity-chromatography stained by coomassie blue (left) or detected by in-gel fluorescence using GFP channel (right). **g** Size-exclusion chromatography of crosslinked NALCN complex. Fractions between red lines were pooled for cryo-EM sample preparation.

**Fig. S2:**
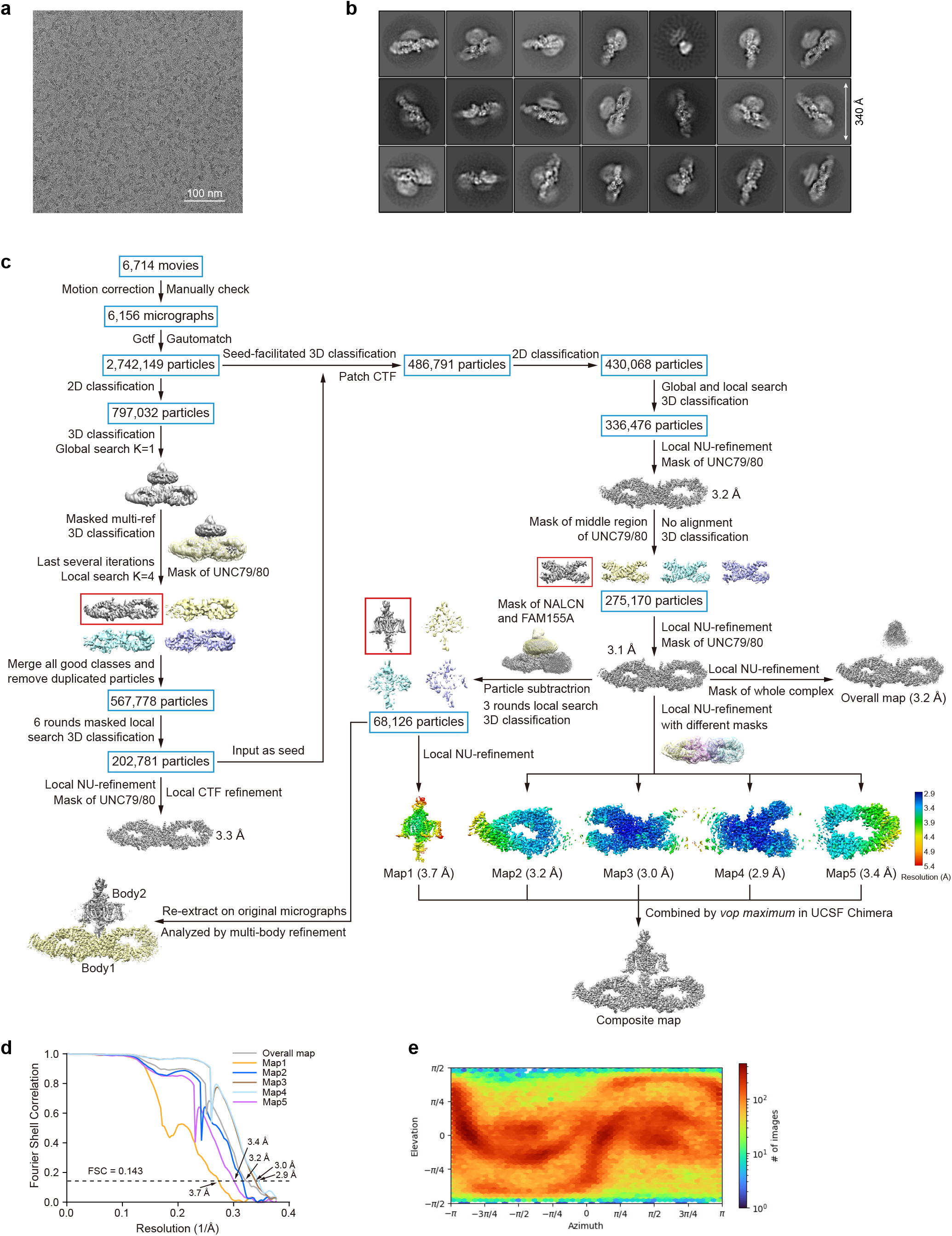
Cryo-EM image analysis of NALCN-FAM155A-UNC79-UNC80 quaternary complex. **a** The raw micrograph of NALCN-FAM155A-UNC79-UNC80 quaternary complex. **b**2D-class averages of NALCN-FAM155A-UNC79-UNC80 quaternary complex with clear features. **c** Cryo-EM data processing workflow. For details, see ‘Cryo-EM image analysis’ in the Methods section. **d** Gold-standard Fourier Shell Correlation (FSC) of focus-refined maps shown in c after correction of masking effects. **e** Angular distribution of consensus refinement of the quaternary complex.

**Fig. S3:**
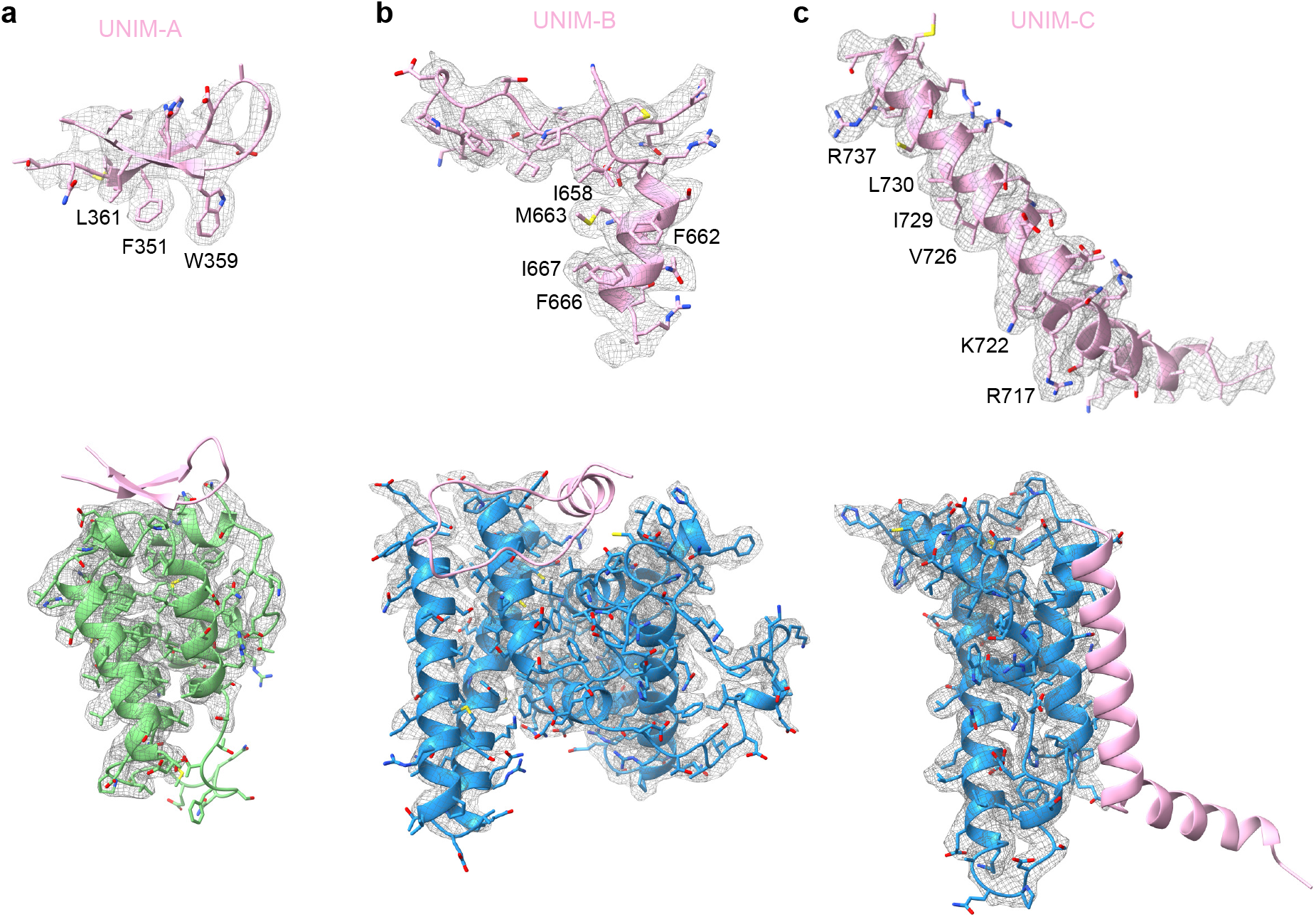
Electron density maps. **a** The electron density map of UNIM-A was shown in gray meshes, with the electron density of its interacting region in UNC79 is shown below. **b** The electron density map of UNIM-B was shown in gray mesh, with the electron density of its interacting region in UNC80 is shown below. **c** The electron density map of UNIM-C was shown in gray mesh, with the electron density of its interacting region in UNC80 is shown below. The contour level was 4-6 σ.

**Fig. S4:**
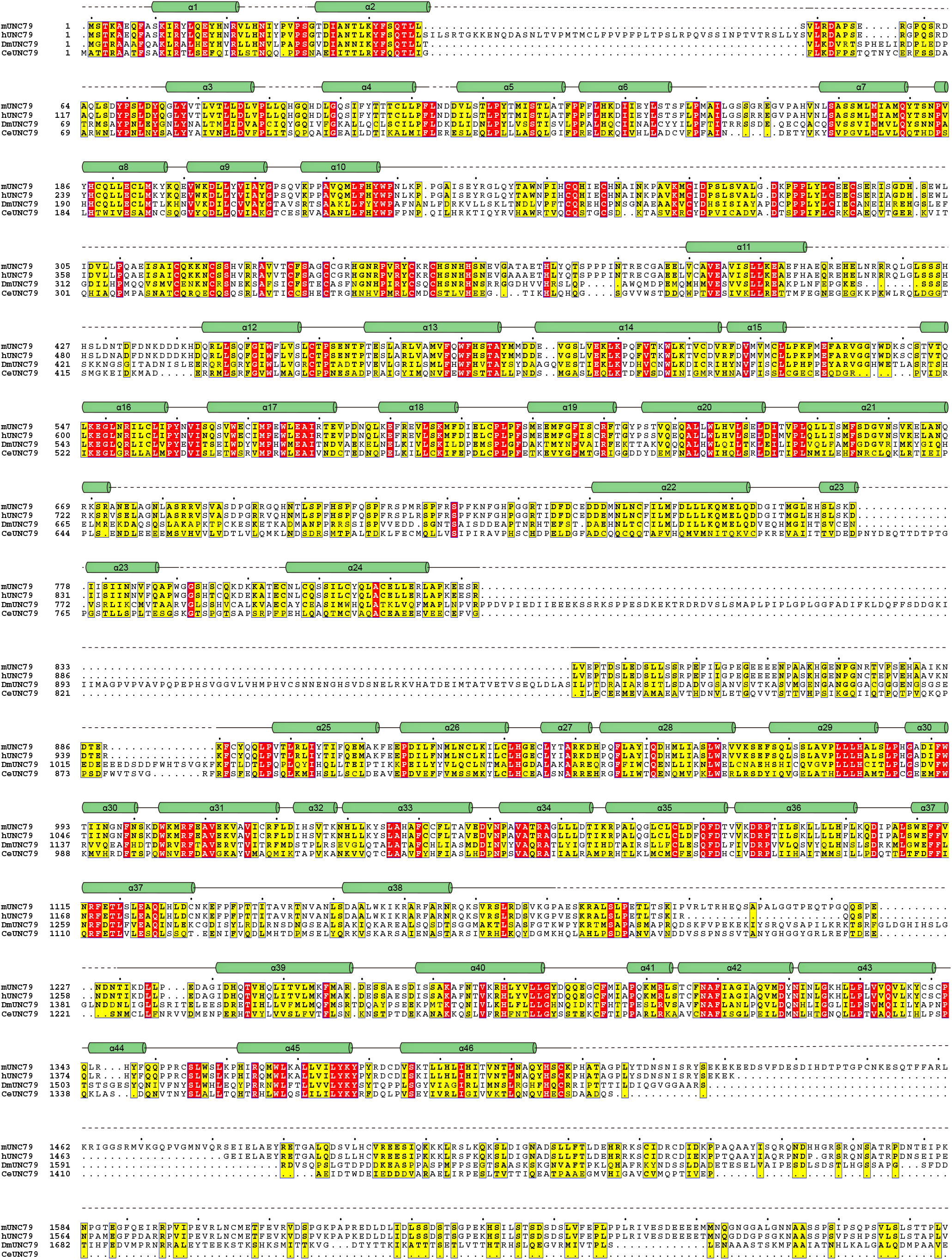
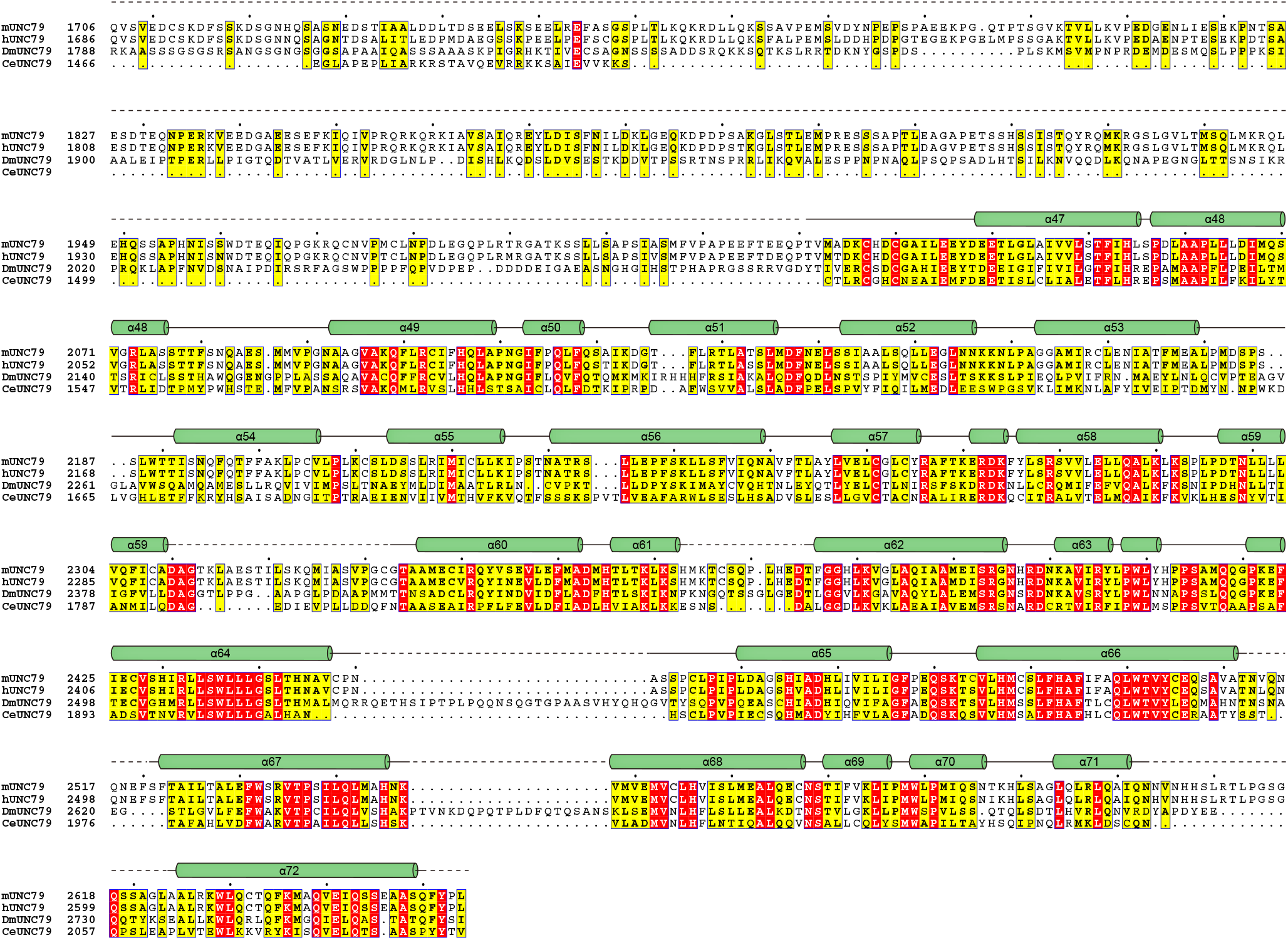
Sequence alignment of UNC79. The sequence alignment of UNC79 from mouse (mUNC79), human (hUNC79), Drosophila melanogaster (DmUNC79), and Caenorhabditis elegans (CeUNC79). Highly conserved and relative conserved residues are shaded in red and yellow, respectively. Secondary structures are shown above and unresolved residues are shown as dashes.

**Fig. S5:**
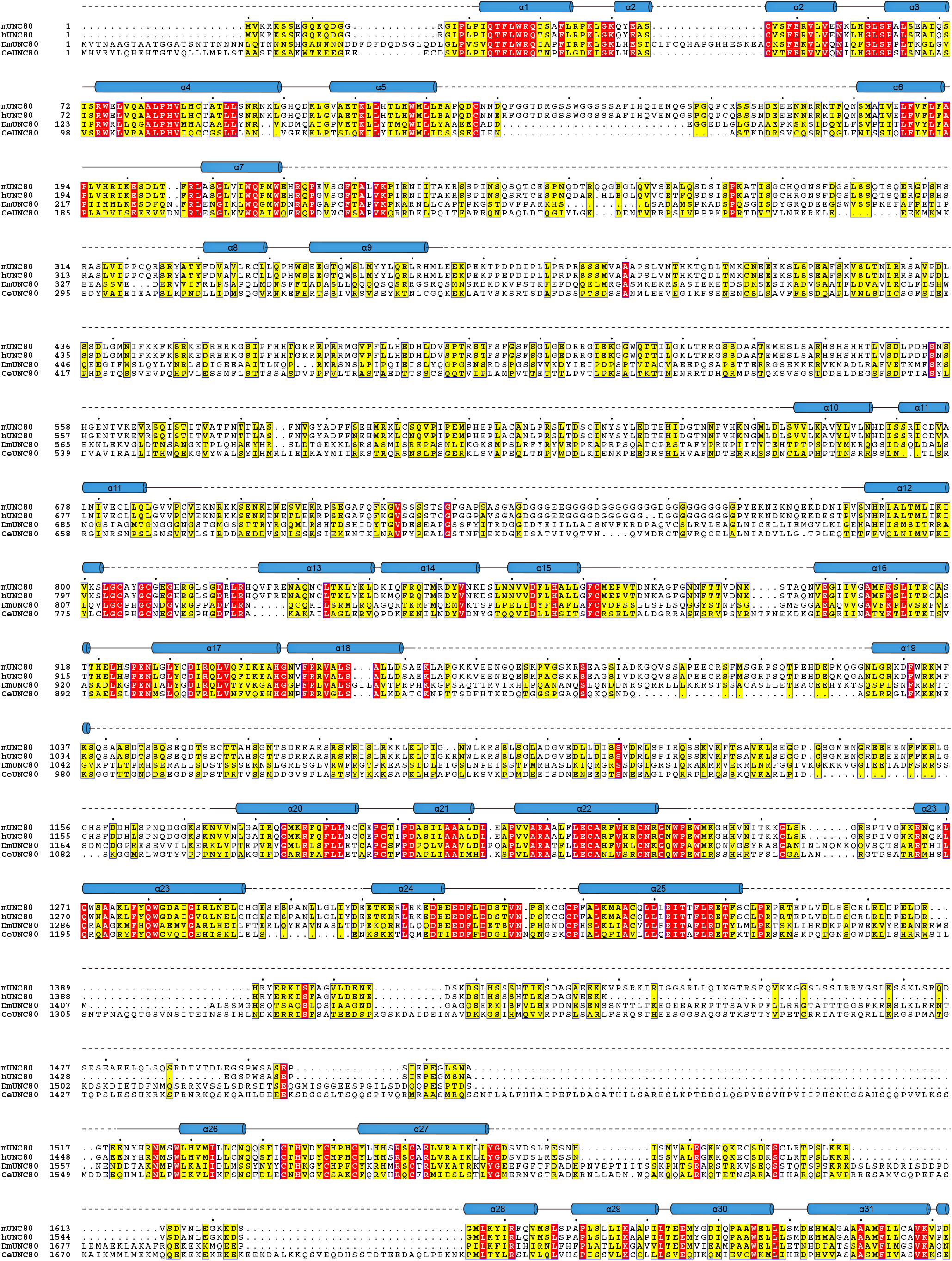
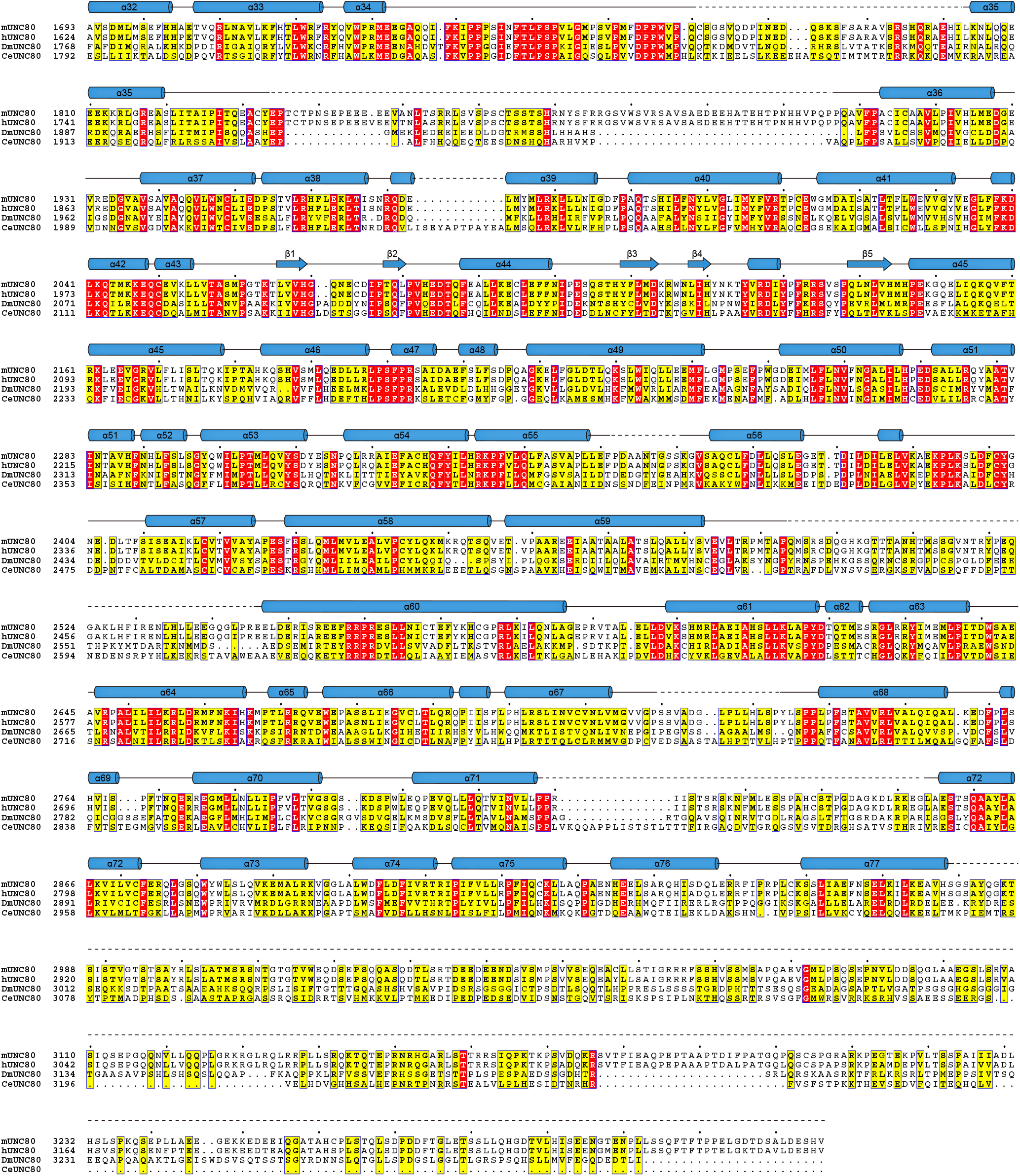
Sequence alignment of UNC80. The sequence alignment of UNC80 from mouse (mUNC80), human (hUNC80), Drosophila melanogaster (DmUNC80), and Caenorhabditis elegans (CeUNC80). Highly conserved and relative conserved residues are shaded in red and yellow, respectively. Secondary structures are shown above and unresolved residues are shown as dashes.

**Fig. S6:**
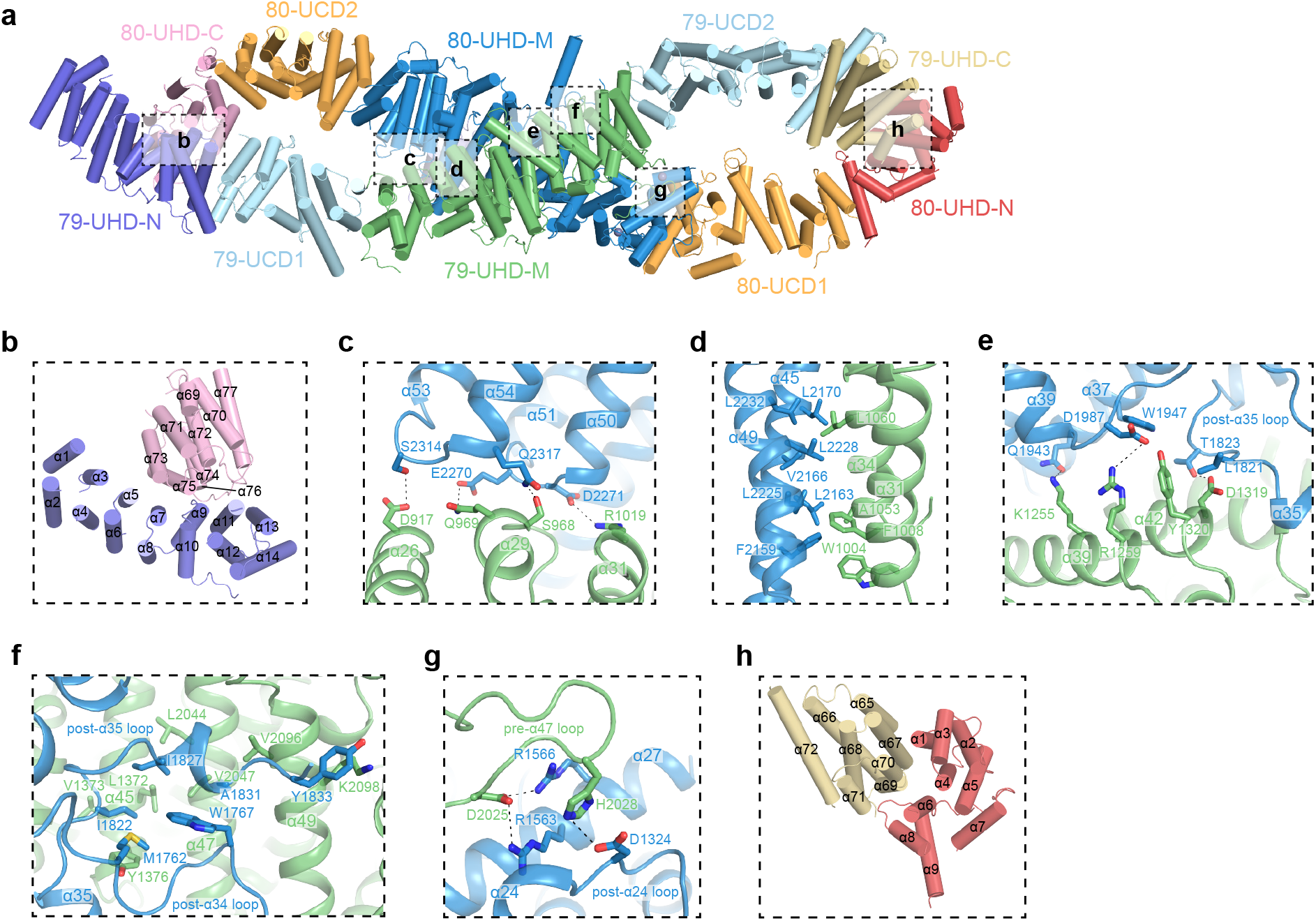
Details of interactions between UNC79 and UNC80. **a** Cartoon representation of UNC79-UNC80 heterodimer. Helices are shown as cylinders. Each domain is colored as in Fig. 2a. **b** The interface between 79- UHD-N and 80-UHD-C boxed in **a**. **c-g** The interface between 79-UHD-M and 80-UHD-M boxed in **a**. **h** The interface between 79-UHD-C and 80-UHD-N boxed in **a**.

**Fig. S7:**
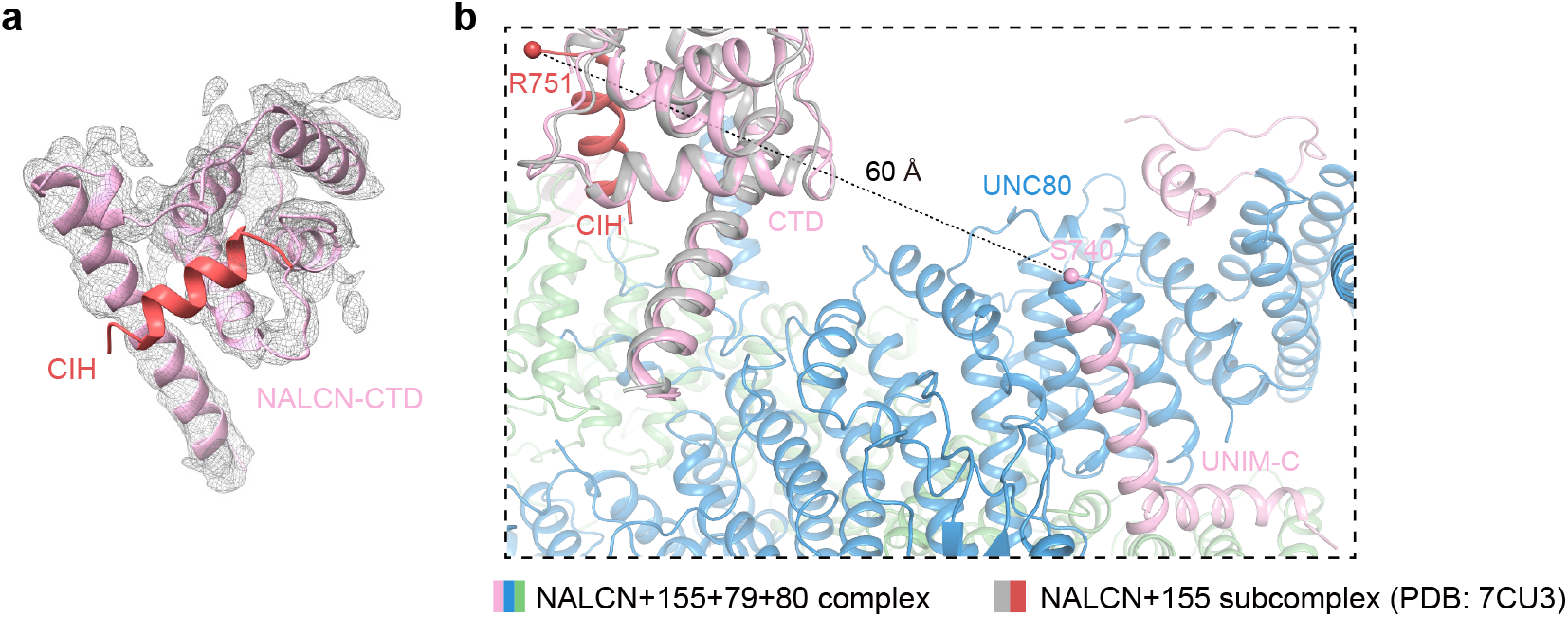
The Structure of NALCN CTD in NALCN-FAM155A-UNC79-UNC80 quaternary complex. **a** Electron density map of NALCN CTD was shown in gray meshes. The density of CIH is absent. The position of CIH (red helix) is based on the structure NALCN-FAM155A subcomplex (PDB ID: 7CU3). The contour level was 4 σ. **b** The structure of the NALCN-FAM155A subcomplex (PDB ID: 7CU3, gray and red) is aligned onto the NALCN-FAM155A-UNC79-UNC80 quaternary complex using the NALCN subunit as reference. The distance between Cα atoms of the N-terminus of CIH (R751) in the NALCN-FAM155A subcomplex and the C-terminus of UNIM-C (S740) is measured and labelled as dashes.

**Table S1:**
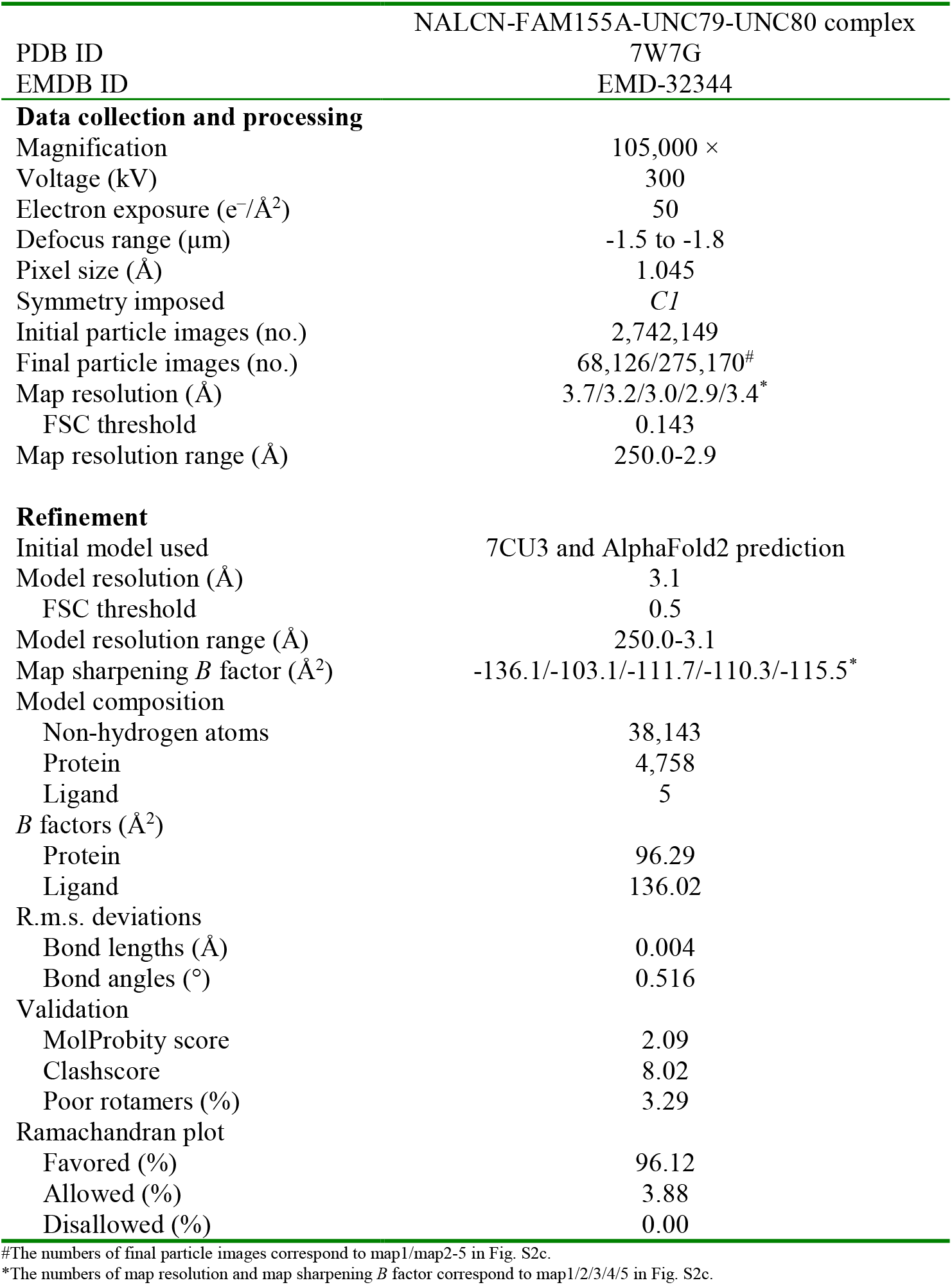
Cryo-EM data collection, refinement and validation statistics.

